# GENOMES UNCOUPLED1: an RNA-binding protein required for early PSII biogenesis

**DOI:** 10.1101/2023.09.21.558905

**Authors:** Chaojun Cui, Shuyi Sun, Siyuan Zhang, Yu Zhang, Chanhong Kim

**Author notes:** **Author contributions:** Conceptualization: CC, CK. Methodology: CC, CK. Investigation: CC, SS, SZ, YZ, CK. Visualization: CC. CK. Funding acquisition: CK. Supervision: CK. Writing – original draft: CC. Writing – review & editing: CK. **Competing interests:** Authors declare that they have no competing interests.

## Abstract

Genomes Uncoupled1 (GUN1), a nuclear-encoded chloroplast pentatricopeptide repeat (PPR) protein, serves as a master integrator of diverse retrograde signals, mediating chloroplast-to-nucleus retrograde signaling. Although PPR proteins primarily function in organelle RNA metabolism, a target RNA of GUN1 remained unknown. This study reveals that GUN1 recognizes *psbD* transcripts derived from a blue light-responsive promoter (BLRP), transcripts referred to as *psbD* BLRP. Overexpression of GUN1 significantly reduces the levels of *psbD* BLRP, whereas the loss of GUN1 leads to the accumulation of the putative target RNA. The *in vitro* RNA and *in vivo* genetic studies further demonstrate the critical role of the C-terminal small MutS-related (SMR) domain in stimulating *psbD* BLRP processing and PsbD (D2, a PSII core protein) synthesis. The GUN1-dependent *psbD* BLRP processing promotes PSII biogenesis during the early seedling development and de-etiolation phase. This finding underscores the role of GUN1 as an RNA-binding protein, highlighting the essential function of the SMR domain in processing the target RNA.

**Significance:** Biogenic retrograde signaling is essential in regulating chloroplast biogenesis, and it entails a nuclear-encoded chloroplast protein, Genomes Uncoupled1 (GUN1). Since its initial discovery in 1993, the *gun1* mutant has been widely used to reveal the precise function of GUN1 in plastids as well as its downstream signaling components. Although GUN1 contains pentatricopeptide repeat motifs, it has been considered a non-canonical PPR protein because no one could demonstrate GUN1-RNA interaction in planta. However, through our computational analysis of PPR codes and subsequent biochemical studies, we were able to unravel its putative target RNA, namely *psbD* BLRP. Therefore, our finding may lead to a reevaluation of GUN1 research in the context of RNA metabolism and a revision of our current understanding of GUN1.

---

Chloroplasts are semiautonomous subcellular organelles whose biogenesis and functions essentially depend on the coordinated expression of nuclear and chloroplast genomes (1–4). The bilateral paths – anterograde (nucleus-to-chloroplast), primarily involved in plastid transcription and translation, and biogenic retrograde (chloroplast-to-nucleus) communications – ensure the accurate stoichiometric assembly of the photosynthetic apparatus and a myriad of chloroplast metabolisms (5, 6). However, once the chloroplast biogenesis becomes constrained, biogenic retrograde signaling overrides the anterograde signaling pathway to repress photosynthesis-associated nuclear genes (PhANGs) through yet unknown molecular mechanisms (7, 8). More than thirty years ago, a forward genetics approach discovered various *genomes uncoupled* (*gun*) mutants with impaired repression of PhANGs to dysfunctional plastids that failed to develop into chloroplasts, opening a new era of retrograde signaling research (7, 9). In particular, *gun1* mutant has been extensively investigated by different laboratories since GUN1 function remains unclear compared to other GUN proteins primarily involved in the tetrapyrrole biosynthesis (TBS) pathways (10–15). Moreover, recent reports about GUN1 added more complexity to our understanding of its function and its role in related retrograde signaling (RS) pathways than was previously thought, as it appears to be involved in multiple chloroplast events, including protein import, RNA editing, proteostasis, and tetrapyrrole biosynthesis (16–19).

GUN1 is a nuclear-encoded chloroplast pentatricopeptide repeat (PPR) protein containing a C-terminal small MutS-related (SMR) domain. While PPR motifs are primarily implicated in RNA metabolisms (20, 21), the SMR domain is thought to play a role in suppressing homologous recombination with an endonuclease activity, as seen in the MutS2 protein, a member of the MutS family (22, 23). However, there has been no report of GUN1-RNA interaction since the initial characterization of the *gun1* mutant by Professor Joanne Chory’s group in 1993 (9). Among those chloroplast-localized PPR-SMR proteins, such as pTAC2 (PLASTID TRANSCRIPTIONALLY ACTIVE 2), SVR7 (SUPPRESSOR OF VARIEGATION 7), SOT1 (SUPPRESSOR OF THYLAKOID FORMATION 1) and GUN1, only the SOT1 protein was shown to interact with plastid-transcribed RNAs and appears to have an endonuclease activity associated with its SMR (24–27). Unlike the *gun1* mutant, *sot1* exhibits a coupled expression of genomes (i.e., the coordinated expression of photosynthesis-associated genes encoded by nucleus and plastid genomes), underlining a unique function of GUN1 in the context of RS (25).

Recent investigations showed that GUN1 associates with distinct groups of chloroplast proteins involved in plastid gene expression (PGE), RNA editing in RNA editosome, proteostasis, or TBS (16, 18, 28, 29). The subsequent reverse genetic studies on *gun1* with a concurrent defect in either plastid proteostasis, PGE, or TBS provided considerable evidence that GUN1 is crucial in regulating the multiplicity of chloroplast homeostasis under various physiological conditions (16, 18, 29). Tadini et al. (2020) also found that GUN1 affects the accumulation of NEP (Nuclear-Encoded Plastid RNA Polymerase)-dependent transcripts and chloroplast protein import in Arabidopsis cotyledons upon perturbation of plastid proteostasis (19). With the complexity of the GUN1 interactome and its roles in a broad spectrum of plastid activities (10), it is rational to support GUN1 as an integrator of multiple retrograde signals emitted from a dysfunctional plastid in varying degrees (10). However, these ranges of GUN1 function were discovered based on the conditions in which exogenous chemical treatments either inhibit plastid proteostasis or carotenoid biosynthesis (7, 10). Therefore, the chemical conditions associated with the earlier GUN1 research call for a reinvestigation of GUN1 under non-invasive conditions. In this regard, the previous findings that the GUN1 protein is relatively abundant in young seedlings, and its absence decelerates cotyledon greening in a proportion of *gun1* siblings, provide a way to explore GUN1 functionality without the need for intrusive chemical treatments (14).

Like other terrestrial plants, the Arabidopsis genome encodes numerous PPR proteins (30). To date, only some of the 441 proteins have been characterized in Arabidopsis. PPR proteins are found to primarily function in organelle RNA metabolisms, including editing, splicing, processing, turnover, and translation. These functions are indispensable for plant development and viability in the ever-changing environment (31–33). Their roles are thought to be RNA sequence-specific, encoded by the PPR codes positioned in each PPR motif through specific di-residues (34–37). Despite extensive investigation and tuning of the decoding algorithm to identify the cognate RNA target sequence, many PPR proteins still have undetermined target RNA sequences.

## GUN1 PPR motifs recognize *psbD* BLRP

As such, this study reexplored the GUN1 PPR code, as we wished that the advancement of PPR research may unveil its putative RNA substrate(s). ScanProsite (https://prosite. expasy.org) predicted 12 PPR motifs with the C-terminal SMR domain in GUN1 (36, 38). We then selected the critical residues required for decoding the PPR code of each PPR motif based on previous reports (36–38), as shown in Figure 1A (see also Fig. S1). This computational approach eventually disclosed potential target nucleotides, which may pair with the PPR motifs of GUN1: the di-residues (6^th^ and 1^st^ residues of the PPR motif) unveiled 11 codes corresponding to 11 potential target nucleotides from the predicted 12 PPR motifs. Combining 6^th^ and 1^st^ di-residues leaves the last PPR motif uncoupled (Fig. S1). We then used the deduced sequence (5’-AA(T>C)(T>C)(C>T)G(T>C)(C>T)GA(C>T)-3’), similar to the recently predicted target sequence (39) except for the affinity-based ambiguity, to find its counterpart in the Arabidopsis plastome. With this approach, we identified several potential target transcripts, including *psbD* BLRP*, rpoC1, atpB*, and *trnL* (Fig. 1*A*). BLRP is referred to as the blue light-responsive promoter activated by the plastid-encoded RNA polymerase and nuclear-encoded plastid sigma factor 5 (SIG5), transcribing extended forms of polycistronic transcripts (called *psbD* BLRP) compared to other BLRP-independent *psbD* transcripts (40–42). A short sequence at the beginning of the 5’-untranslated region (UTR) in *psbD* BLRP is predicted to be decoded by GUN1 PPR motifs, as indicated in Figure 1B. To explore whether GUN1 affects the metabolism of any of these potential target transcripts, we first examined their abundances using Northern blot analyses with target-specific probes (Fig. 1*B*).

**Fig. 1.**
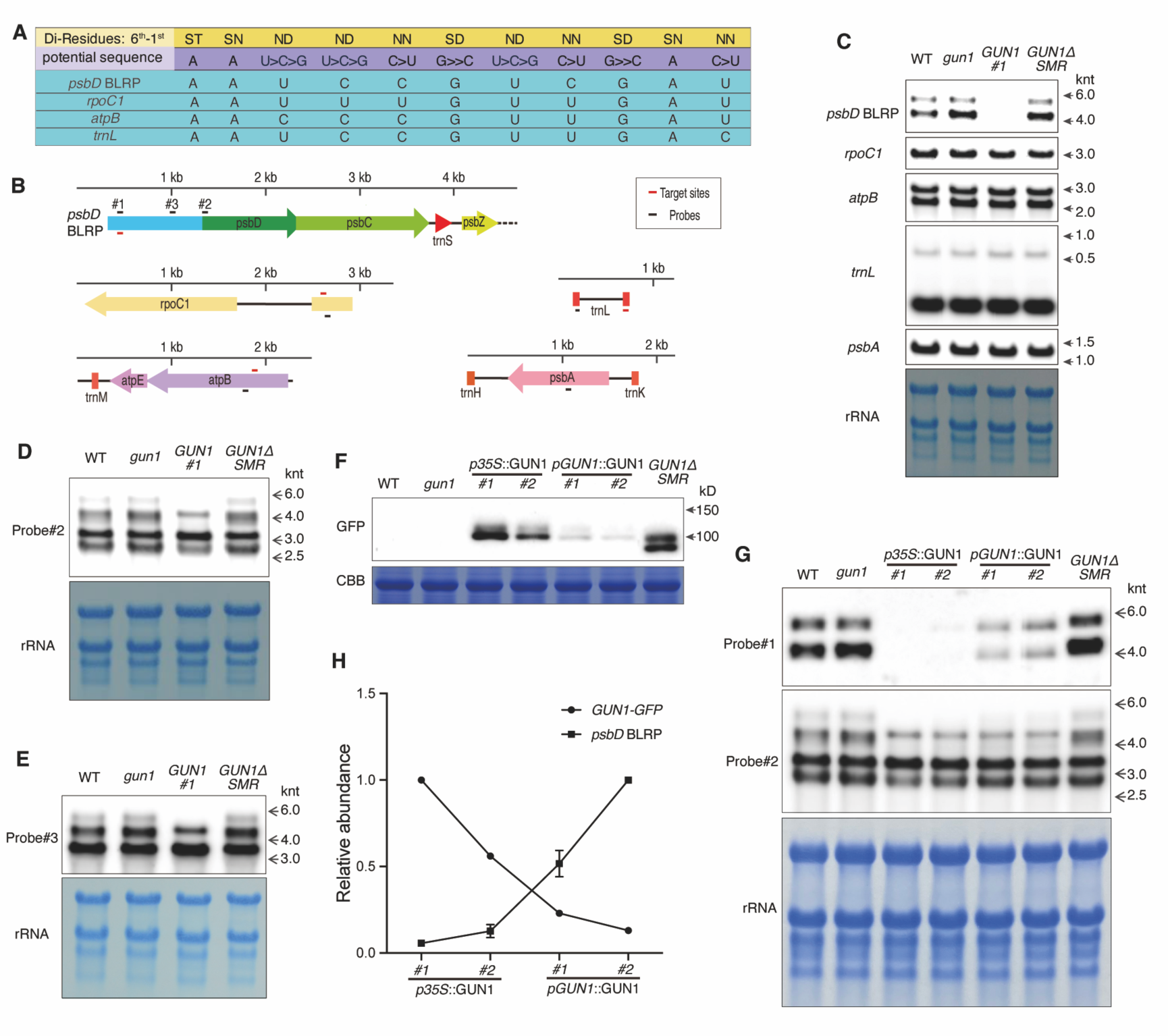
GUN1 decodes and processes the *psbD* BLRP transcripts. (*A*) The predicted target RNA sequences of GUN1 PPR motifs. The first row shows the ‘di-residues (6^th^-1^st^)’ in each PPR motif whose combinatorial configuration is critical in predicting potential target nucleotides (second row). The ambiguous sequence was BLASTed, and the most related target sequences in the chloroplast genome are shown in the rows below, including *psbD* BLRP, *rpoC1*, *atpB*, and *trnL*. (*B*) The schematic representation of the predicted target nucleotides in each transcript (red bars). The probe positions for Northern blot analyses are indicated (black bars). *psbA* encoding PSII D1 protein is shown as an unrelated control. (*C*) RNA gel blot analyses with total RNA isolated from 5-day-old seedlings of WT, *gun1*, *gun1 p35S::GUN1-GFP* (*GUN1#1*), and *p35S::GUN1△SMR-GFP* (*GUN1△SMR*) grown under non-stress conditions. Probe#1 was used for the detection of the *psbD* BLRP transcripts. The positions of RNA size markers are indicated (kilo nucleotide, knt) at the right. The methylene blue staining of rRNAs is used as the loading control. (*D*-*E*) Northern Blot analyses were conducted with probes #2 and #3, as shown in (B). (*F*) The Western blot result shows the GUN1-GFP or GUN1ΔSMR-GFP fusion protein levels in independent *gun1* transgenic lines. The protein size corresponding to 100 kilodaltons (kD) is indicated. Coomassie brilliant blue (CBB) staining of RbcL (Ribulose bisphosphate carboxylase large chain) is used as the loading control. (*G*) Northern blot analysis for the independent transgenic lines (F) demonstrates the different levels of the *psbD* transcripts. (*H*) The quantification of the GUN1-GFP and the *psbD* BLRP transcripts abundances in four transgenic lines (F). The band intensity was determined by ImageJ 1.53k. The protein band intensity of each genotype was compared to that in *p35S::GUN1#1*. The transcript band intensity of each genotype was compared to that in *pGUN1::GUN1#2*. Data represent the mean of three biological replicates with a cognate standard error. Western and Northern blot analyses were repeated using three independent biological replicates with reproducible results.

## GUN1 processes *psbD* BLRP in an SMR-dependent manner

Total RNA was prepared from 5-day-old seedlings and resulting RNA blot analyses detected the steady-state transcript levels of the potential targets (Fig. 1*C*). In addition to WT and *gun1*, we also included total RNAs extracted from the transgenic *gun1* seedlings expressing either the cauliflower mosaic virus 35S promoter (p35S)-driven GUN1-GFP (GUN1) or the SMR domain deleted GUN1△SMR-GFP (GUN1△SMR). Note that confocal imaging showed the foci-like GFP signal of the GUN1 fusion protein in chloroplasts, but it showed aggregated signals in the case of GUN1△SMR (Fig. S2*A* and *B*). The GFP signal of both fusion proteins gradually declined with seedling growth (Fig. S2*C* and *D*). Therefore, we decided to use young seedlings, where GUN1 is shown to be stable, to determine the levels of target transcripts. To detect *psbD* BLRP transcripts in young seedlings, we first used a 5’ UTR probe#1 (−888 to −856 from the start codon of *psbD*) including the target region of the GUN1 PPR motif. The subsequent RNA blot analyses identified two bands, one around 4.3-knt and one below 6.0-knt. Remarkably, both signals disappeared in the GUN1 overexpression line with a complete rescue of a *gun1* phenotype (i.e., the deficiency of cotyledon greening in a proportion of seedlings; Fig. S3) (14), while they accumulated in *gun1* and *gun1 GUN1△SMR* relative to WT seedlings. Surprisingly, the *psbD* coding region probe (*psbD* CR probe#2, +1 to +33 from the *psbD* start codon) developed a band around 4.3-knt in the GUN1 overexpression line along with those BLRP-independent transcripts (detected in all genotypes examined) (Fig. 1*B* and *D*). We thus presumed that probe#2-detected ∼4.3-knt transcript(s) in the GUN1 overexpression line might be a processed form of *psbD* BLRP transcripts, lacking at least the 5’ UTR target region. The target RNA alteration seems likely to be mediated by the SMR domain as manifested by the accumulation of probe#1-hybridized *psbD* BLRP transcripts in both *gun1* and *gun1 GUN1ΔSMR* seedlings. To ensure that the *psbD* CR probe#2-detected ∼4.3-knt RNA in the GUN1 overexpression line is a processed *psbD* BLRP, we designed another 5’ UTR probe#3 (−243 to −214 from the *psbD* start codon) to carry out RNA gel blot analysis (Fig. 1*B*). We thought that if the transcript detected by probe#2 is derived from initial *psbD* BLRP transcripts, 5’ UTR probe#3 will also detect transcript(s) at ∼4.3-knt in the GUN1 overexpression line (Fig. 1*B* and *E*). The RNA blotting result indeed confirmed the emergence of the processed *psbD* BLRP in the GUN1 overexpression line. This notion also suggests that ∼4.3-knt *psbD* BLRP detected by probe#1 might undergo similar processing by GUN1 in the presence of GUN1 target sequence. Indeed, the intensity of the RNA band between 4-knt and 3-knt in the GUN1 overexpression line was higher than in other genotypes (Fig. 1*B, D* and *E*). Furthermore, the significant inverse correlation between the *psbD* BLRP transcript level and GUN1 protein abundance was observed in several independent *gun1* transgenic seedlings expressing *GUN1-GFP* under the control of either p35S or native promoter (pGUN1), corroborating *psbD* BLRP as a plausible target of GUN1 (Fig. 1*F*-*H*). The transcript levels of the other potential targets of GUN1 PPR, such as *rpoC1*, *atpB*, and *trnL*, and the unrelated transcript *psbA* remained unchanged (Fig. 1*C*).

## GUN1 physically interacts with *psbD* BLRP to promote early chloroplast biogenesis

Next, we wondered whether the abnormal accumulation of *psbD* BLRP transcripts decelerates D2 synthesis and chloroplast biogenesis during early *gun1* seedling development (14). If this hypothesis holds true, the decelerated D2 synthesis during early chloroplast biogenesis could explain the appearance of seedlings with white cotyledons, dependent on the threshold level of D2. When D2 levels surpass the threshold, chloroplast biogenesis occurs, while levels below the threshold may slow cotyledon greening (Fig. S3). As such, the correlation between *psbD* BLRP accumulation and D2 synthesis (or PSII activity) was investigated during de-etiolation, where the transition of the etioplast to the chloroplast and the light-dependent biogenesis of the photosynthesis apparatus are readily monitored with great precision. To evade any unknown impacts of GUN1 overexpression, we selected a *gun1* transgenic line expressing native promoter-driven GUN1-GFP, whose expression was confirmed by Western blot analysis (Fig. S4). Although the exogenous *GUN1* transcript level was slightly higher than endogenous *GUN1* (Fig. S4), we selected the stable transgenic line because of its lowest *GUN1-GFP* transcript level compared to other independent transgenic lines. A *gun1 GUN1△SMR* line was included as a control. RNA gel blot analysis and the subsequent immunoblotting found an opposite correlation between the level of *psbD* BLRP and D2 abundance (Fig. 2*A*-*C*). The persistence of *psbD* BLRP, especially upon illumination in both *gun1* and *gun1 GUN1△SMR-GFP*, led to a reduction in D2 protein levels, resulting in a significantly decreased maximum quantum yield of PSII photochemistry (F_v_/F_m_) (Fig. 2*D*). No significant change of D1 was observed (Fig. 2*B*). Since the chlorophyll fluorescence parameter F_v_/F_m_ reflects the efficiency of absorbed light energy utilization in photosynthesis (43), the lower F_v_/F_m_ value observed in *gun1* seedlings compared to others suggests a significantly decelerated photosynthetic activity and/or chloroplast biogenesis in *gun1* during de-etiolation. A similar role of GUN1 was also observed in light-grown seedlings (Fig. S5). Significantly, RNA immunoprecipitation (RIP) followed by qPCR analyses confirmed a direct interaction between GUN1-GFP and *psbD* BLRP (Fig. 2*E*). However, the *psbD* BLRP was barely detected in the RIP sample of transgenic plants overexpressing chloroplast-targeted free GFP (cpGFP; Fig. 2*E*). To investigate the interaction between GUN1 and *psbD* BLRP transcripts *in vitro*, we conducted an electrophoretic mobility shift assay (EMSA) using GUN1 protein (132-918 aa) with a 6×His tag, which was expressed in mammalian Expi293 cells. Meanwhile, a 20-nucleotide natural RNA sequence (RNA1) containing the target sequence of GUN1 was synthesized. Two additional RNA sequences, RNA2 and RNA3, were used as control, with 5-nucleotide and 10-nucleotide mutations in the GUN1 target region, respectively (Fig. 2*F*). The EMSA results indicate that only RNA1 showed a shift after incubation with GUN1 protein, while RNA2 and RNA3 did not (Fig. 2*G*). This suggests that the GUN1 protein can target the target RNA region of the *psbD* BLRP transcripts *in vitro*. The results indicate that GUN1 physically interacts with *psbD* BLRP transcripts. All these results imply that GUN1-dependent *psbD* BLRP processing is required to promote D2 synthesis during early chloroplast biogenesis.

**Fig. 2.**
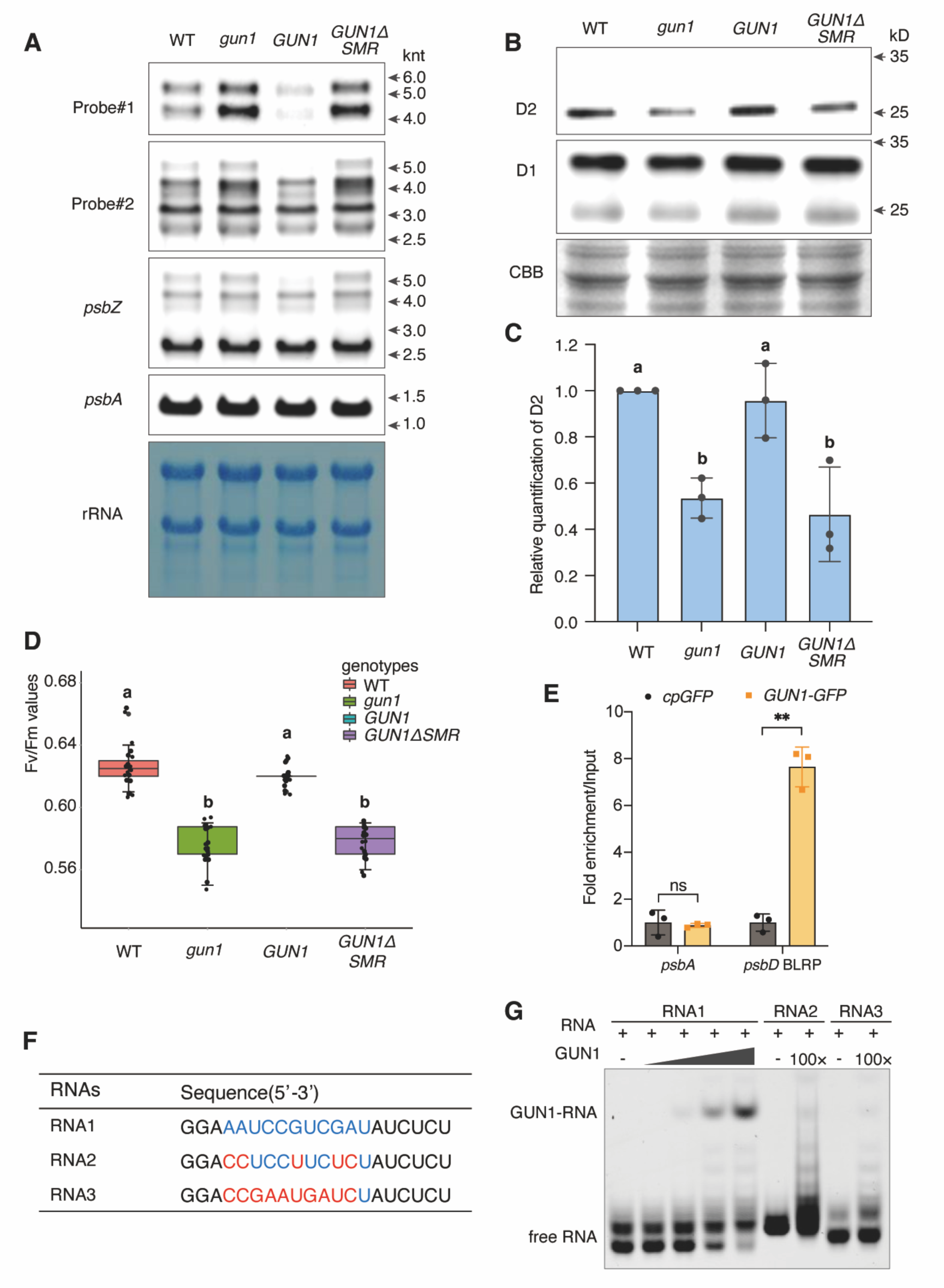
GUN1 promotes D2 synthesis and PSII biogenesis upon the onset of de-etiolation. (*A*) Northern blot analysis of *psbD* BLRP, *psbD*, *psbZ*, and *psbA* in 2.5-day-old etiolated seedlings exposed to light for 6 hours (h). For *psbZ*, the gene-specific probe was used, as shown in Table S2. The rRNAs are shown as the loading control. *pGUN1::GUN1-GFP* (*GUN1*) was selected based on its expression level of GUN1 protein, as shown in Fig S4 (*GUN1#32*). The same growth condition was used for (B-D). (*B*) Western blot analyses reveal the corresponding D1 and D2 levels. CBB staining of RbcL was used as the loading control of protein samples. (*C*) The immunoblotting of the D2 protein was repeated with three independent biological samples with reproducible results. The target band was quantified, and values represent the means ± SD (*n* = 3 replicates). The lowercase letters indicate statistically significant differences between mean values (*P* < 0.05, one-way ANOVA with Tukey’s post hoc honestly significant difference [HSD] test) (*D*) F_v_/F_m_ values. At least 10 seedlings of each genotype (A) were used to measure F_v_/F_m_. The value represents the means ± SD (*n* = 10). This experiment was repeated with three independent replicates with similar results. The lowercase letters indicate statistically significant differences between mean values (*P* < 0.01, one-way ANOVA with Tukey’s post hoc honestly significant difference [HSD] test). (*E*) RNA-immunoprecipitation (RIP) coupled with RT-qPCR analyses using *GUN1-GFP* (*GUN1*) and *gun1* expressing *cpGFP* (signal peptide of Ribulose bisphosphate carboxylase small subunit fused with GFP) after the onset of de-etiolation. RNAs were co-immunoprecipitated using GFP-trap beads, and the input and eluate were used for RT-qPCR analysis to examine the interaction between GUN1 and *psbD* BLRP. The enrichment value was normalized to the input, representing means ± SD (*n* = 3). Asterisks indicate statistically significant differences relative to the control (*P* < 0.01, Student’s *t*-test). ns: not significant. (*F*) The RNA sequences used for EMAS analysis: RNA1 contained the original sequence of the *psbD* BLRP transcripts with the target sequence (blue) of GUN1 protein. RNA2 was designed as a control containing five mutation sites (red) within the GUN1 target region. RNA3 had an additional five mutation sites in the GUN1 target region, based on the RNA2 sequence. The RNAs were labeled with Cy5 (cyanine fluorescent dye) at the 3’ ends. (*G*) The EMSA analysis was conducted to examine the interaction between GUN1 and RNA *in vitro*. The reaction mixture (10 µL) contained 10 nM RNA and varying concentrations of the truncated GUN1 (132-918 aa; 0, 15.625, 62.5, 250, 1000[100×] nM).

## SMR domain of GUN1 plays a critical role in *psbD* BLRP processing

Because the SMR domain of GUN1 retains critical amino acid residues in the motifs (LDxH and TGxG, where x represents any amino acid; Fig. 3*A*) necessitated for endonuclease activity (23), we hypothesized that SMR activity might be indispensable for *psbD* BLRP processing. As proof of concept, we mutated those residues to monitor how they influence the *psbD* BLRP abundance. The chosen substitutions in the LDxH motif were Leu(L)784-to-Ala(A), D785-to-A, and His(H)787-to-A, respectively. For the TGxG motif, the substitutions were Thr(T)823-to-A, Gly(G)824-to-A, and G826-to-A, respectively (Fig. 3*A*). To reduce the possibility of the tag interfering with endonuclease activity, we used a small Myc (∼1kDa) tag instead of the GFP (∼27kDa). All constructs were stably transformed *in planta* to generate various transgenic lines in the *gun1* mutant background. As a control, we also created stable *gun1 GUN1-Myc* transgenic lines (Fig. 3*B*). Concerning whether any of those conserved residues in the SMR domain play a critical role in chloroplast biogenesis, we monitored the phenotype of 5-day-old transgenic seedlings. Except for L784A, the substitution of either D785A, H787A, T823A, G824A, or G826A failed to rescue the *gun1*-conferred decelerated cotyledon greening phenotype (Fig. 3*C*). The levels of *psbD* BLRP transcripts were also monitored in those transgenic seedlings during de-etiolation. The RNA gel blot analysis revealed that also here, except for L784A, either substitution, D785A, H787A, T823A, G824A, or G826A, failed to prevent the *psbD* BLRP accumulation (Fig. 3*D*). These results indicate that the endonuclease activity of the SMR domain is necessary to facilitate *psbD* BLRP processing and is vital for early chloroplast biogenesis upon germination.

**Fig. 3.**
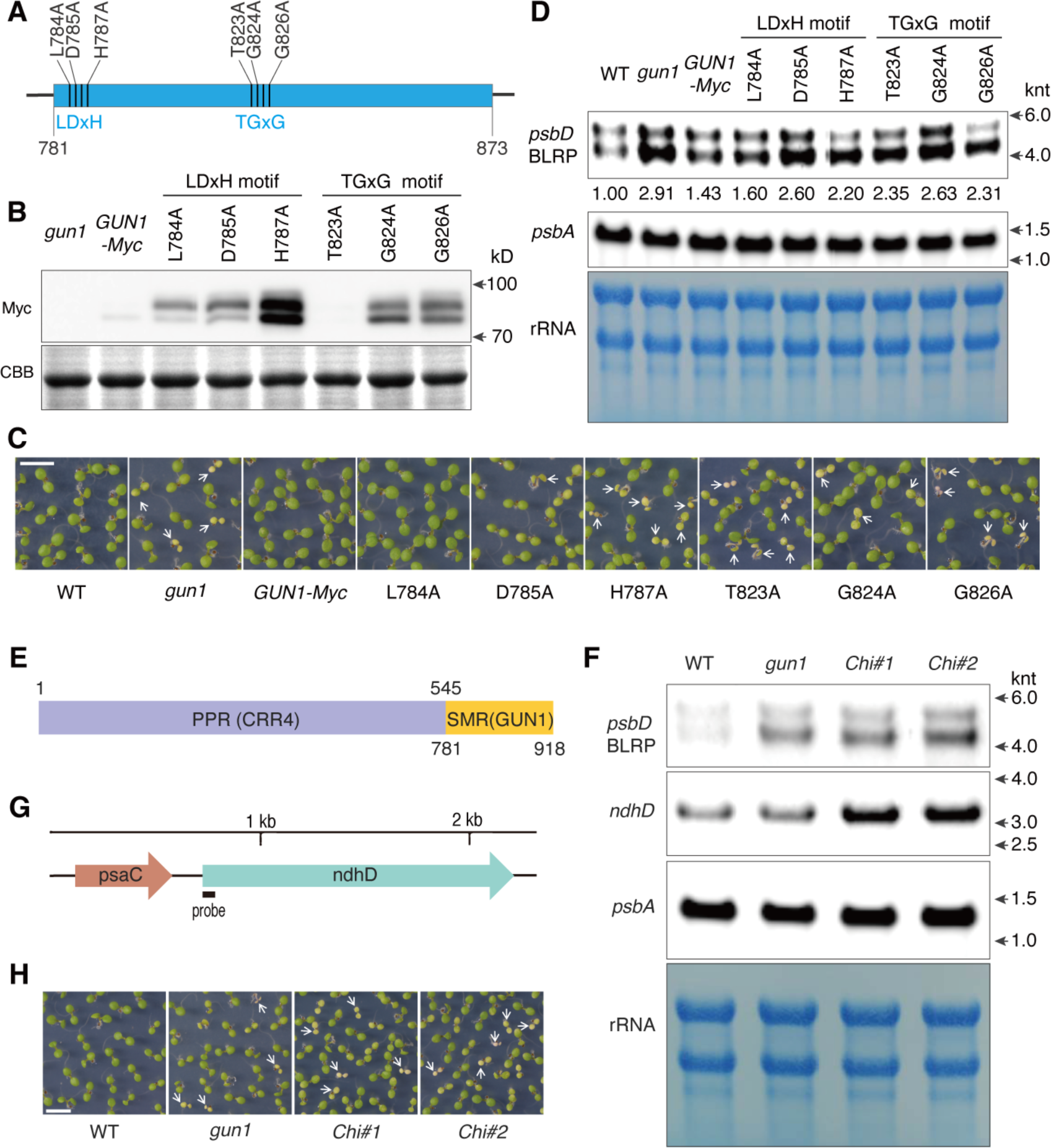
Both SMR domain and PPR motifs of GUN1 are indispensable to process *psbD* BLRP transcripts. (*A*) LDxH and TGxG motifs in the SMR domain were previously shown to be essential in the endonuclease activity of SOT1 (23, 24). Each critical amino acid of two motifs was mutated into Ala, and GUN1 variants fused with the Myc tag were stably expressed in *gun1* for complementation assay. (*B*) Western blot result shows the levels of Myc-tagged GUN1 variants in the stable *gun1* transgenic lines. One of each representative line was chosen for Western blot analysis among multiple independent lines with comparable phenotypes. CBB staining of RbcL is shown as the loading control. (*C*) The *gun1*-caused emergence of a group of seedlings with white or pale cotyledons was monitored in light-grown 5-day-old seedlings. The white arrows mark the pale or bleached seedlings. Scale bar, 5 mm. (*D*) Northern blot analysis. During de-etiolation, the steady-state level of *psbD* BLRP was analyzed in the indicated genotypes using probe#1. *psbA* was used as a control. The numbers below each band indicate the intensity of *psbD* BLRP transcripts normalized based on the intensity of the *psbA* transcript. The ImageJ 1.53k was used for quantification. The methylene blue staining of rRNA is used as the loading control. (*E*) Chimeric GUN1 protein. PPR motifs of CRR4 (CRR4PPR) and SMR domain of GUN1 (GUN1SMR) are fused. (*F*) Northern blot analysis. Two independent transgenic lines of *gun1 CRR4PPR-GUN1SMR* (*Chi#1* and *Chi#2*) were included along with WT and *gun1*. *psbA* was used as a control. The steady-state levels of *psbD* BLRP (probe#1), *ndhD*, and *psbA* were examined after the onset of de-etiolation. The methylene blue staining of rRNA was used as the loading control. This experiment was repeated using three independent biological replicates for each genotype with reproducible results. (*G*) The probe’s position is shown. (*H*) The images are representative phenotypes of light-grown 5-day-old seedlings. The white arrows indicate the pale or bleached seedlings. Scale bar = 5 mm.

## PPR motifs of GUN1 are indispensable for *psbD* BLPR recognition

Conversely, we also explored the specificity of GUN1 PPR motifs in decrypting its target RNA by creating a chimeric PPR-SMR protein. To replace GUN1 PPR with functionally unrelated PPR, we chose CRR4, whose PPR motifs are required to edit the start codon of the *ndhD* transcript in chloroplasts (44). The construct of CRR4 PPR motifs fused with the SMR of GUN1 was stably transformed into *gun1* (Fig. 3*E*). The independent transgenic lines (*Chi#1* and *Chi#2*; Fig. S6) were examined for their *psbD* BLRP phenotype using Northern blot analysis. The result showed similar or slightly higher levels of *psbD* BLRP in the transgenic plants than *gun1* (Fig. 3*F*), indicating that the GUN1 PPR code is specific for *psbD* BLRP and that the GUN1 SMR activity requires GUN1 PPR motifs to process *psbD* BLRP. It was noticeable that the potential RNA target of CRR4, *ndhD* was upregulated in both transgenic lines relative to WT (Fig. 3*G* and *F*). Perhaps, CRR4 also plays a role in stabilizing *ndhD* in chloroplasts. Besides, the chimeric protein failed to rescue the *gun1*-conferred cotyledon phenotype (Fig. 3*H*).

## An anterograde path from CRY1 to GUN1 modulates early chloroplast biogenesis

A plastid-encoded bacterial-type RNA polymerase (PEP) recognizes the BLRP, in which nuclear-encoded chloroplast-localized SIGMA FACTOR 5 (SIG5) is known to facilitate target promoter recognition (45, 46). Similar to white light, blue light rapidly induces the expression of *SIG5*, and knockdown of *SIG5* considerably impairs the expression of *psbD* BLRP (45). A nuclear cryptochrome called blue light receptor CRY1 is involved in activating *psbD* BLRP (47), so *SIG5* is considered a downstream target of CRY1. We then hypothesized that the inactivation of CRY1 might revert the accumulation of *psbD* BLRP in *gun1*. However, despite the loss of CRY1 in *gun1*, the defect of D2 accumulation upon the onset of de-etiolation might remain unchanged if the GUN1-dependent processing of *psbD* BRLP, albeit less expressed, promotes D2 synthesis. If this notion is valid, the data will allow us to depict a novel anterograde pathway from CRY1 to GUN1, promoting *psbD* BRLP-dependent D2 synthesis and chloroplast biogenesis (Fig. 4*A*). To this end, we created *cry1 gun1* double mutants and noticed that loss of CRY1 seemingly reverted the *psbD* BLRP phenotype in *gun1*, but neither the D2 protein level nor the de-etiolation defect (Fig. 4*B*-*D*).

**Fig. 4.**
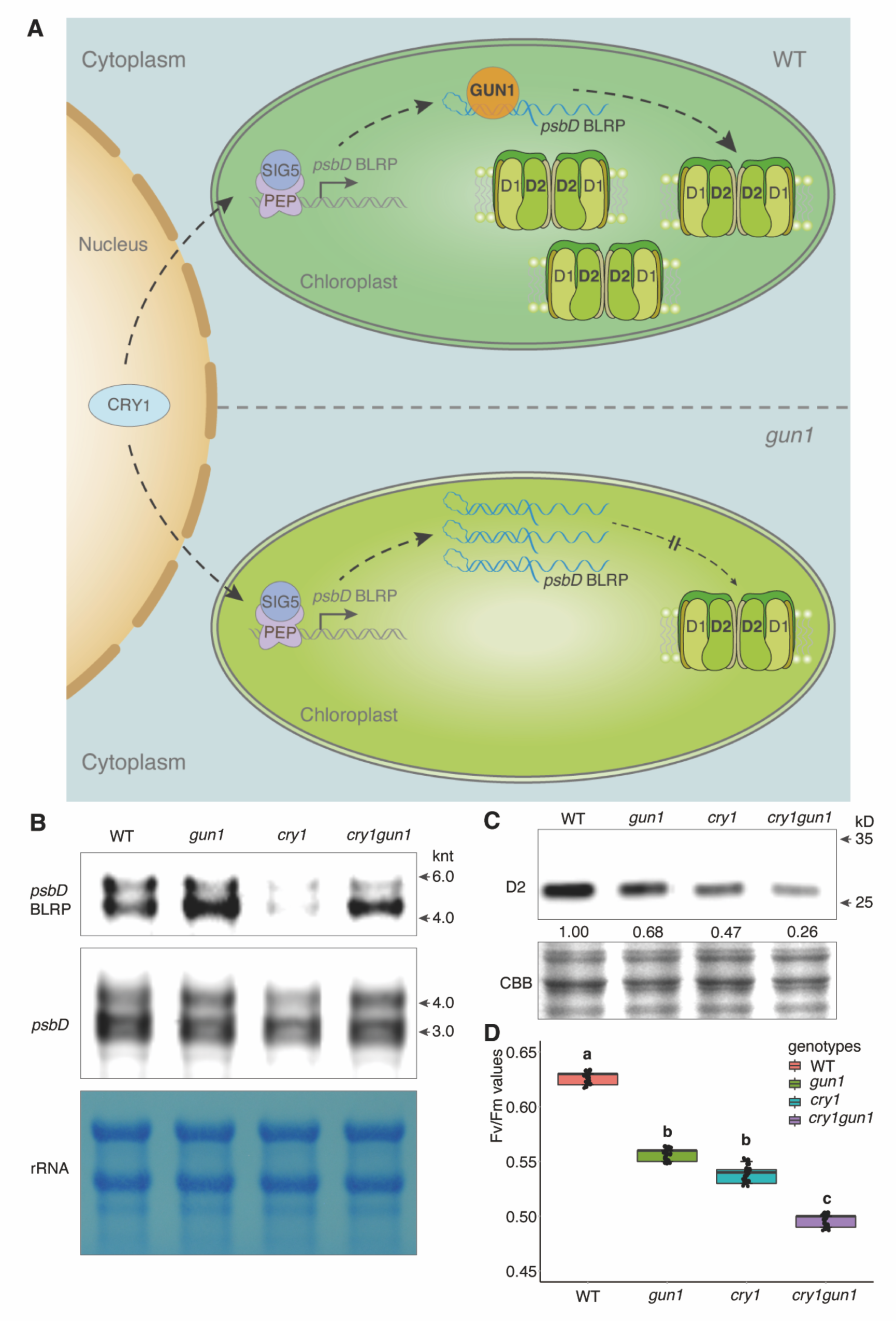
The anterograde pathway from CRY1 to GUN1 promotes early PSII biogenesis. (*A*) Working model showing an anterograde pathway from CRY1 to GUN1, enabling *psbD* BLRP-dependent D2 synthesis and PSII biogenesis. The blue light receptor CRY1 positively regulates the expression of *psbD* BLRP through SIG5, leading to GUN1-dependent processing of *psbD* BLRP to contribute to PSII biogenesis and phototropic growth of young seedlings. (*B*) *cry1* mutation is epistatic to *gun1* mutation in the level of *psbD* BLRP transcripts. The representative result shows the *psbD* BLRP (probe#1) and *psbD* (probe#2) levels in WT, *gun1*, *cry1*, and *cry1gun1* after the de-etiolation onset. (*C*) Western blot result shows the corresponding D2 protein levels of (B). Numbers below the bands indicate the relative intensity of D2 protein levels compared to WT. The band intensity was determined using ImageJ 1.53k. (B) and (C) were repeated in triplicate with reproducible results with independent biological replicates. (*D*) The F_v_/F_m_ value of (B). Values represent the means ± SD (*n* = 12). The lowercase letters indicate statistically significant differences between mean values (*P* < 0.01, one-way ANOVA with Tukey’s post hoc honestly significant difference [HSD] test).

In conclusion, the present study demonstrated that the PPR motif and SMR domain of GUN1 decode *psbD* BLRP and promote its processing, facilitating PSII biogenesis during early seedling development. Given the emergence of the target site-excluding *psbD* transcript(s) using different probes (Fig. 1*D* and *E*), it is conceivable that GUN1 might cleave or process the target RNA to promote D2 synthesis. However, in this case, other PPR or RNA-binding proteins may protect the processed RNA from degradation. Nonetheless, unveiling the direct physical interaction between GUN1 and *psbD* BLRP further raises the question of whether the steady-state levels of *psbD* BLRP or D2 (or PSII activity) are directly associated with the activation of GUN1-mediated retrograde signaling. Certainly, we cannot exclude the possibility that *psbD* BLRP is one of the GUN1 substrates. More plastidial RNAs may interact with GUN1 for its multifunctionality for other biological processes, including protein import.

Moreover, a recent functional conservation study of GUN1 underlined that GUN1 is an ancient protein primarily involved in chloroplast gene expression, while its role in chloroplast retrograde signaling likely evolved more recently (39). Therefore, investigating the co-evolution of the GUN1 PPR code and its target RNA(s) may enable us to investigate potential causes for the emergence of retrograde signaling.

## Materials and Methods

### Plant materials and growth conditions

Wild-type (WT) and mutants used in this study were all in Arabidopsis (*Arabidopsis thaliana*) Col-0 ecotype background. All seeds were harvested from plants grown in the soil (PINDSTRUP) under continuous light conditions (CL, 100 µmol.m^−2.^s^−1^ at 22 ± 2℃). The T-DNA insertion mutant of *gun1* (*SAIL_290_D09*) and *cry1* (*SALK_042397C*) were obtained from the Nottingham Arabidopsis Stock Centre (NASC). T-DNA insertion and knockout of the target gene were confirmed by means of PCR-based genotyping. Seeds were surface-sterilized with 20% hypochlorite solution for 5 min, washed 4-5 times in sterilized water, and then plated on Murashige and Skoog (MS) agar medium (Duchefa Biochemie) supplemented with 0.5% (w/v) sucrose, pH 5.75. After a 2-day stratification at 4℃ in the dark, seeds germinated and grew under CL conditions. *Nicotiana benthamiana* plants are grown under long days conditions (LD, 100 µmol. m^−2^.s^−1^ at 22 ± 2℃, 16-h light/8-h dark cycle). Four-week-old *N. benthamiana* plants were used for all transient expression assays. For the de-etiolation experiment, the etiolated seedlings initially grown under continuous dark conditions for 2.5-day were placed in a light chamber (100 µmol.m^−2.^s^−1^ at 22 ± 2℃) for 6 h prior to harvesting. The shoots were collected to check the protein and RNA expression after freezing with liquid Nitrogen.

### Vector construction and generation of transgenic plants

Each cDNA corresponding to GUN1 (918 aa) and GUN1△SMR (780 aa), lacking the stop codon, was cloned into a pDONR221 Gateway vector through Gateway BP reaction (Invitrogen). After sequencing verification, the entry vector was subsequently recombined into the Gateway-compatible plant binary vectors pGWB505(48) for C-terminal fusion with EGFP (Enhanced Green Fluorescent Protein) through Gateway LR reaction (Invitrogen). The 1053 bp native promoter, fused with the stop codon-less coding sequence of GUN1, was cloned into pGWB504 to generate the native promoter-controlled GUN1-GFP construct. Each GUN1 variant, with a substitution in either amino acid in LDxH and TGxG motifs, and the SMR domain-deleted GUN1 (without stop codon), were subcloned into pGWB517 destination vectors for the C-terminal fusion with a 4×MYC tag. As a control construct, the signal peptide of RBCS was fused with free GFP. Subsequently, all constructs were destined for the binary vector pGWB505. The pGWB505 vectors with each gene of interest were transformed into *Agrobacterium tumefaciens* strain GV3101. The primer sequences used for constructions are shown in Table S1. The floral dip procedure (49) generated Arabidopsis transgenic plants via Agrobacterium-mediated transformation. Homozygous transgenic plants were selected on the MS medium containing 25 mg/L Hygromycin (Thermo Fisher Scientific). For transient expression, Agrobacterium suspensions carrying different constructs were injected into healthy leaves of 4-week-old *N. benthamiana* plants, and the target protein was analyzed 36-48 h later.

### Prediction of GUN1-binding sequences

The protein sequence of GUN1 was obtained from The Arabidopsis Information Resource (TAIR, https://www.arabidopsis.org) and scanned in the ScanProsite (https://prosite.expasy.org/) to predict PPR motifs. The prediction revealed twelve PPR repeats in the GUN1 protein. We then selected key di-residues in each PPR motif (sixth and first residues) to encrypt the PPR codes of GUN1 (37). Based on the previous studies on PPR codes (36–38), the di-residues of GUN1 PPR motifs unveiled ambiguous target nucleotide sequence: 5’-AA(TC)(TC)(CT)G(TC) (CT)GA(CT)-3’. Then, the complete chloroplast genome was downloaded from National Center for Biotechnology Information (https://www.ncbi.nlm.nih.gov/) to search for the target transcript position in SnapGene (https://www.snapgene.com). Searching with ambiguous sequences allowed us to find potential target transcripts (Fig. S1).

### RNA extraction and qRT–PCR

Total RNA was extracted using the Universal Plant Total RNA Extraction Kit (BioTeke) and spectrophotometrically quantified on the NanoDrop 2000 (Thermo Fisher Scientific). One µg of total RNA was reverse-transcribed using the HiScript II Q RT SuperMix (Vazyme Biotech) according to the manufacturer’s instruction. The RT-qPCR was performed using the ChamQ Universal qPCR Master Mix (Vazyme Biotech) on the QuantStudioTM 6 Flex Real-Time PCR System (Applied Biosystems). The relative transcript level was calculated by the ΔΔCT comparative quantification method (50) and normalized to the housekeeping gene *ACTIN2* (AT3G18780). The primer sequences for qRT-PCR are shown in Table S1.

### RNA gel blot analysis

Total RNA was extracted using the TRIzol Reagent^TM^ solution (Thermo Fisher Scientific). In addition to 2 µl RNA marker (Thermo Fisher Scientific), 3 µg RNA from each sample was used to mix with 2×RNA loading dye (Thermo Fisher Scientific), denatured at 65℃ for 10 min, and placed on ice for 3 min. Samples were loaded into 1% formaldehyde agarose gels and separated at 3V/cm volt for 2 h in 1× RNA running buffer (1× MOPS and 6%[v/v] formaldehyde). Subsequently, the RNA was blotted onto the Hybond^TM^-N^+^ nylon transfer membrane (GE Healthcare) by soaking in 20× SSC (3.0 M NaCl and 0.3 M sodium citrate) overnight, crosslinked with a CX-2000 UV Crosslinker at 1,000 uJ.CM^−2^ × 100 for 15 sec, then hybridized in 10 mL hybridization buffer (Sigma-Aldrich) containing 20 µM. L^−1^ of 3’-biotin-labeled primer probe purified by high-pressure liquid chromatography (HPLC) for 4-12 h at 42°C in Hybaid Shake ‘n’ Stack (Thermo Fisher Scientific) with gentle rotation. The probe sequences are listed in Table S1. After the incubation, the membrane was washed twice for 5 min at 42°C with high salt wash buffer (2×SSC, 0.1% [w/v] SDS) and washed twice for 10 min at 42°C with low salt wash buffer (0.1× SSC, 0.1% [w/v] SDS). According to the manufacturer’s instructions, the hybridization between probe and RNA was detected using a Chemiluminescent Nucleic Acid Detection Module kit (Thermo Fisher Scientific).

### Protein extraction and western blot analyses

Total proteins were extracted from seedlings using the homogenization buffer (0.0625 M Tris-HCl [pH 6.8], 1% [w/v] SDS, 10% [v/v] glycerin, and 0.01% [v/v] 2-mercaptoethanol) and then quantified with the Pierce BCA protein assay kit (Thermo Fisher Scientific). The total proteins were separated by 8%-12% SDS-PAGE gels and blotted onto an Immune-Blot polyvinylidene difluoride (PVDF) membrane (Bio-Rad). All the antibodies were obtained from commercial suppliers (anti-GFP and anti-MYC antibodies from Roche; anti-D2 antibody from Agrisera). Coomassie brilliant blue (CBB) R250 staining for RbcL was used as a loading control.

### Determination of photochemical efficiency

The maximum photochemical efficiency of PSII for a single plant or a population of young seedlings was determined into F_v_/F_m_ values in a FluorCam system (FC800-C/1010GFP; Photon Systems Instruments) according to the instrument manufacturer’s instructions. For the light-grown young seedlings, 3-to 6-day-old seedlings were selected individually to measure the F_v_/F_m_ values. The F_v_/F_m_ values were determined in de-etiolating seedlings after 6 h of light illumination for the de-etiolation experiment.

### RNA immunoprecipitation assay

The aerial part of 6 h-de-etiolated seedlings was collected, frozen in liquid nitrogen, and ground to a fine powder. Approximately the powder corresponding to 1.0 g fresh weight samples was lysed using cold 4*(w/v) RIPA buffer (10 mM Tris-HCl [pH 7.5], 150 mM NaCl, 0.5 mM EDTA, 0.1% [w/v] SDS, 0.5% [w/v] sodium deoxycholate, 1% [v/v] Triton X-100, 1 mM PMSF, 1× Complete protease inhibitor cocktail [MedChemExpress], and 4 U/ml Recombinant RNasin® Ribonuclease inhibitor [Promega]) on ice with 10-15 brief vortexes within 15 min. The lysed samples were kept on ice for 10 min. After centrifuging twice at 12,500 *g* for 10 min at 4°C, approximately 100 µl of the supernatants were taken as input samples, and the remaining supernatant (∼2.7 ml) was incubated with 30 µL of GFP-Trap magnetic agarose beads (GFP-TrapMA; Chromotek) for 2 h at 4°C by gentle vertical rotation. The magnetic agarose beads were washed twice using RIPA buffer for 5 min at 4°C by gently vertical rotation. After incubation, the beads were washed 2-5 times with RIPA buffer. The RNA that was bound to target protein and RNA from input samples were purified from the beads with TRI-Reagent^TM^ solution (Thermo Fisher Scientific), precipitated by adding 1 volume (∼300 µl) of isopropanol, 7 µl glycogen (5 mg/ml), and 5 µl of 3 M sodium acetate (pH 5.2) for overnight at −80°C, and washed with 1 ml 75% (v/v) ethanol. The RNA from beads was eluted with 12 µl RNase-free water, and the RNA from input was dissolved in 20 µl RNase-free water. The total RNA from beads and 1 µg RNA from input were used as templates to synthesize cognate complementary DNA via reverse transcriptases (Vazyme Biotech). RT-qPCR was then carried out with the specific primer sets for *psbD* BLRP and *psbA* (used as a negative control), listed in Table S1. The RIPA buffer was prepared with DEPC (diethyl pyrocarbonate)-treated H_2_O.

### EMSA assay

The coding sequence of the GUN1 (132–918) was amplified from *A. thaliana* cDNA by PCR and subsequently cloned into a modified pMlink vector with a protein A tag at the N-terminus. The plasmid was transfected to Expi293 cells using polyethylenimine (PEI) when the cells reached a density of 2.5×10^6^/mL. After approximately 60 h of culture following transfection, the cells were harvested and then lysed in lysis buffer containing 50 mM Tris-HCl pH 7.7, 300 mM NaCl, 10% (v/v) glycerol, 5 mM MgCl_2_, 0.5 mM EDTA, 0.25% CHAPS, 5 mM ATP, 2 mM DTT, 1 µg/mL Aprotinin, 1 µg/mL Pepstatin, and 1 µg/mL Leupeptin at 4 ℃ for 40 min. The lysate was centrifuged at 15,000 g for 1 h at 4℃. The supernatant was incubated with IgG beads (Smart-Lifesciences) for 1 h, and the beads were washed five times (10 mL each) with wash buffer containing 50 mM Tris-HCl pH 7.7, 300 mM NaCl, 10% (v/v) glycerol, 5 mM MgCl_2_, 0.5 mM EDTA, 0.1% CHAPS, 5 mM ATP, 2 mM DTT. The tags were removed by on-column cleavage for 1 h, and the eluted protein was dialyzed in storage buffer containing 50 mM Tris-HCl pH 7.7, 100 mM NaCl, 10% (v/v) glycerol, 2 mM DTT. Finally, GUN1 (132-918 aa) was concentrated to ∼0.7 mg/mL and stored at −80°C. The target RNAs were synthesized with Cy5 labeling at 3’end. RNA1 (5’-GGAAAUCCGUCGAUAUCUCU-3’) is the part natural sequence in the UTR region of the *psbD* BLRP transcripts with the predicted target sequence. RNA2 (5’-GGACCUCCUUCUCUAUCUCU-3’) is a control RNA with 5 mutation sites. RNA3 (5’-GGACCGAAUGAUCUAUCUCU-3’) is the other control RNA with 10 mutation sites. The Protein and RNA reaction mixtures were incubated at 37°C for 30 min in the EMSA buffer (50 mM Tris-HCl pH 7.7, 100 mM NaCl, 10% [v/v] glycerol, 2 mM DTT), and subsequently loaded on 5% native polyacrylamide gels. The RNA was visualized by fluorescence imaging.

### Confocal laser-scanning microscopy

The sub-plastidial localization of GFP-fused proteins, GUN1-GFP and GUN1ΔSMR-GFP, were examined in both *N. benthamiana* leaves (following transient expression) and stable Arabidopsis transgenic lines under confocal laser-scanning microscopy (TCS SP8, Leica Microsystems). GFP fluorescence was excited with an argon laser at a wavelength of 488 nm, and emission was detected with a 500–530 nm filter. Chlorophyll fluorescence was obtained between 650 and 700 nm. All the images were obtained using Leica LAS AF Lite software, version 2.6.3 (Leica Microsystems).

### Accession Numbers

To find the sequence information of genes explored in this study, you may check the following accession numbers in the Arabidopsis Information Resource (https://www.arabidopsis.org). *GUN1* (At2g31400), *CRR4* (At2g45350), *SIG5* (At5g24120), *CRY1* (At4g08920), *psbD* (AtCg00270), *rpoC1* (AtCg00180), *atpB* (AtCg00480), *trnL* (AtCg00400), *psbA* (AtCg00020), and *psbZ* (AtCg00300).

## Data and materials availability

All data are available in the main text or the supplementary materials.

## Acknowledgments

We thank Dr. Yanfei Mao for the technical help of RNA gel blot analyses; and the core facility of Cell Biology for their assistance in confocal microscopy. We thank Dr. Keun Pyo Lee, Dr. Joanna Melonek, and Prof. Ian Small for their insightful feedback. We dedicate this study to Prof. Joanne Chory.

---

**Figure S1.**
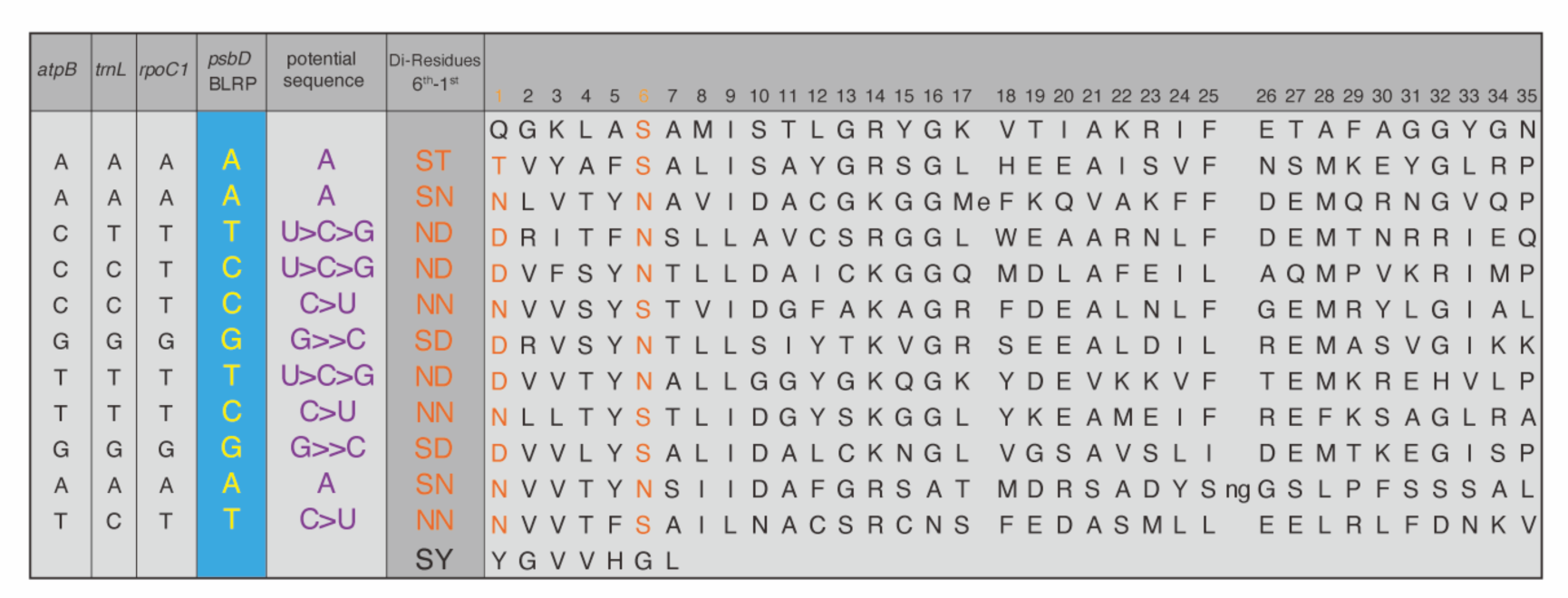
Predicted nucleotides encoded by GUN1 PPR motifs. ScanProsite (https://prosite. expasy.org/) revealed 12 probable PPR motifs. The key di-residues in each PPR motif are shown in orange, and the potential target nucleotides are indicated in purple. The ambiguous sequences were closely matched with specific regions of four transcripts expressed in chloroplasts, i.e., *psbD* BLRP, *rpoC1*, *trnL*, and *atpB,* respectively.

**Figure S2.**
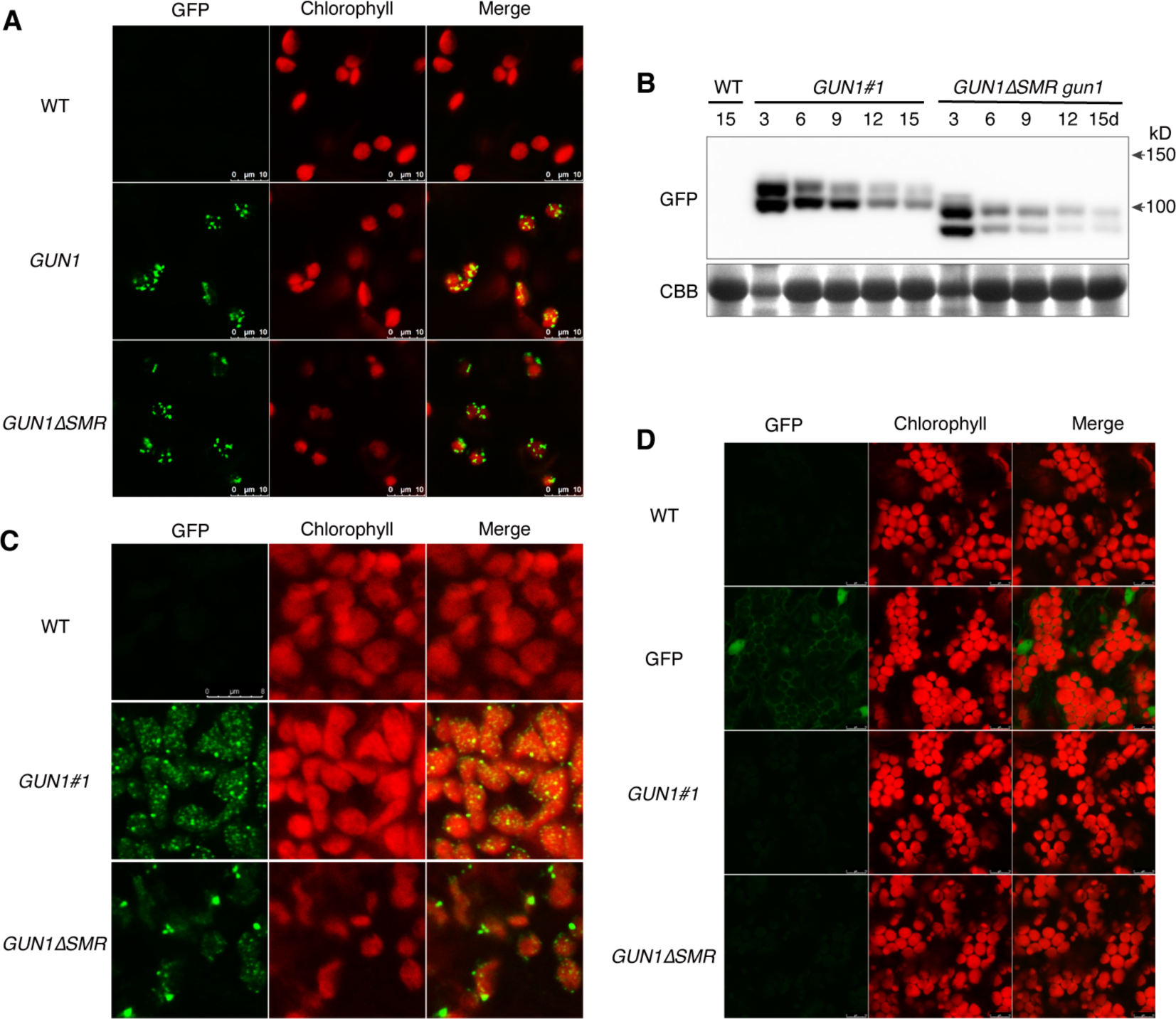
Subcellular localization of GFP-tagged GUN1 and GUN1ΔSMR. (*A*) The confocal images represent the GFP signal of GUN1 and GUN1ΔSMR in *Nicotiana benthamiana* leaves. The 35S promoter-driven construct was each transiently expressed through Agrobacterium infiltration, and the GFP signal was monitored at 48 h following infection by confocal microscopy. (*B*) The confocal images represent the GFP signal of GUN1 and GUN1ΔSMR in chloroplasts of light-grown 3-day-old *gun1* transgenic seedlings. (*C*) The relative GUN1 protein levels during seedling development. CBB staining of RbcL is used as the loading control. The result turned out to be reproducible with independent biological replicates. (*D*) The GFP signal of GUN1 and GUN1ΔSMR in cotyledons of 10-day-old seedlings. Then, the transgenic WT seedling expressing free GFP was used as technical control.

**Figure S3.**
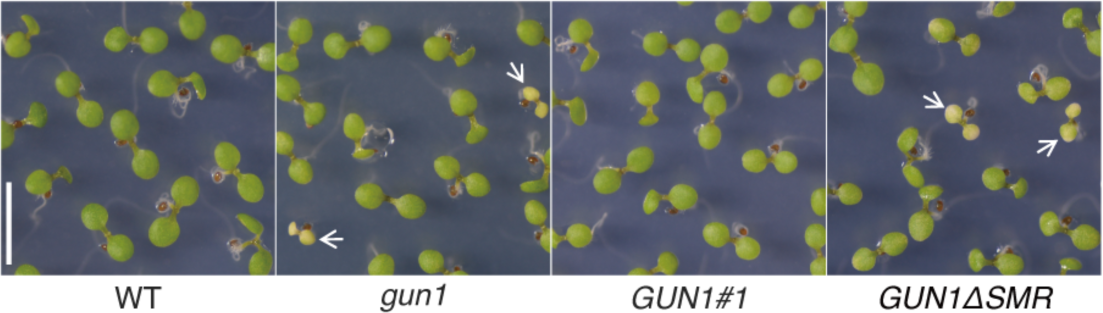
The SMR domain is required to rescue the albino cotyledon phenotype of *gun1*. The representative images of 5-day-old WT, *gun1*, *GUN1#1*, and *GUN1ΔSMR* seedlings are shown. All seedlings were grown under continuous light conditions (100 ± *µ*mol m^−2^ s^−1^, 22 °C ± 2 °C).

**Figure S4.**
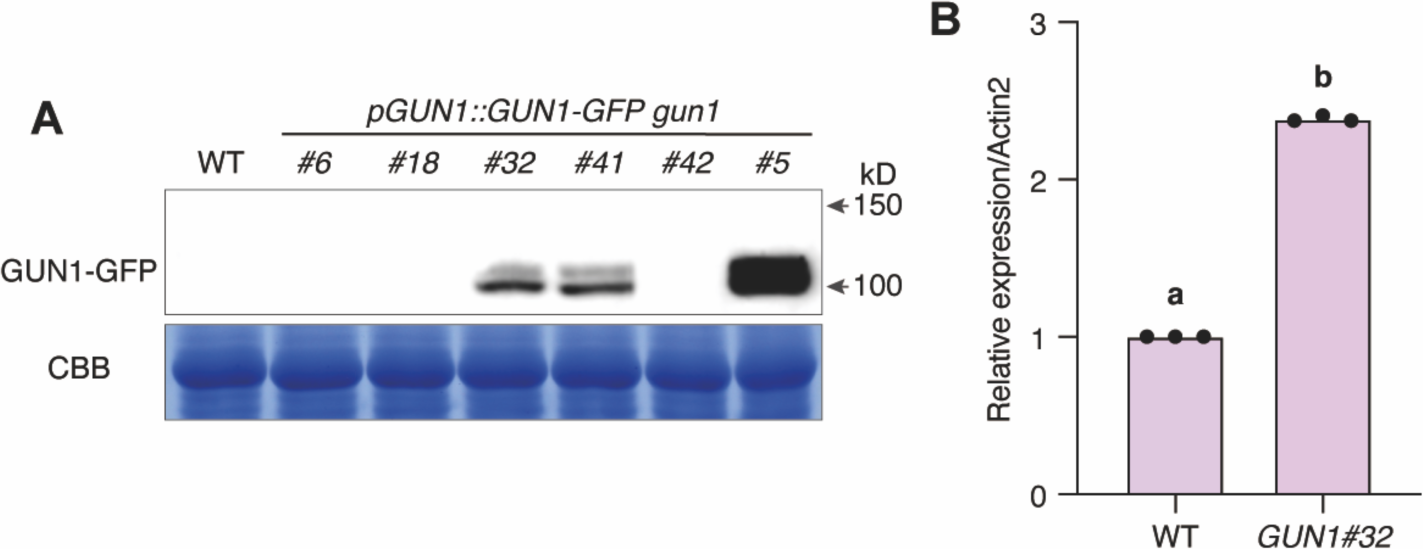
Selection of a *pGUN1::GUN1-GFP gun1* transgenic line. (*A*) Six independent *gun1* transgenic lines expressing GUN1-GFP under the control of the native promoter (pGUN1) were examined to determine the relative abundance of GUN1-GFP using Western blot analysis. (*B*) Line #32 (A) was further selected to determine the cognate transcript level relative to endogenous GUN1 in WT seedlings. For the Western and qRT-PCR analysis, light-grown 5-day-old seedlings were used. The line, *pGUN1::GUN1-GFP gun1#32* (*GUN1)* was used for the de-etiolation experiments. The different lowercase letters indicate statistically significant differences between mean values. *P*-value was obtained from two-tailed Student’s *t*-tests. *P* < 0.01.

**Figure S5.**
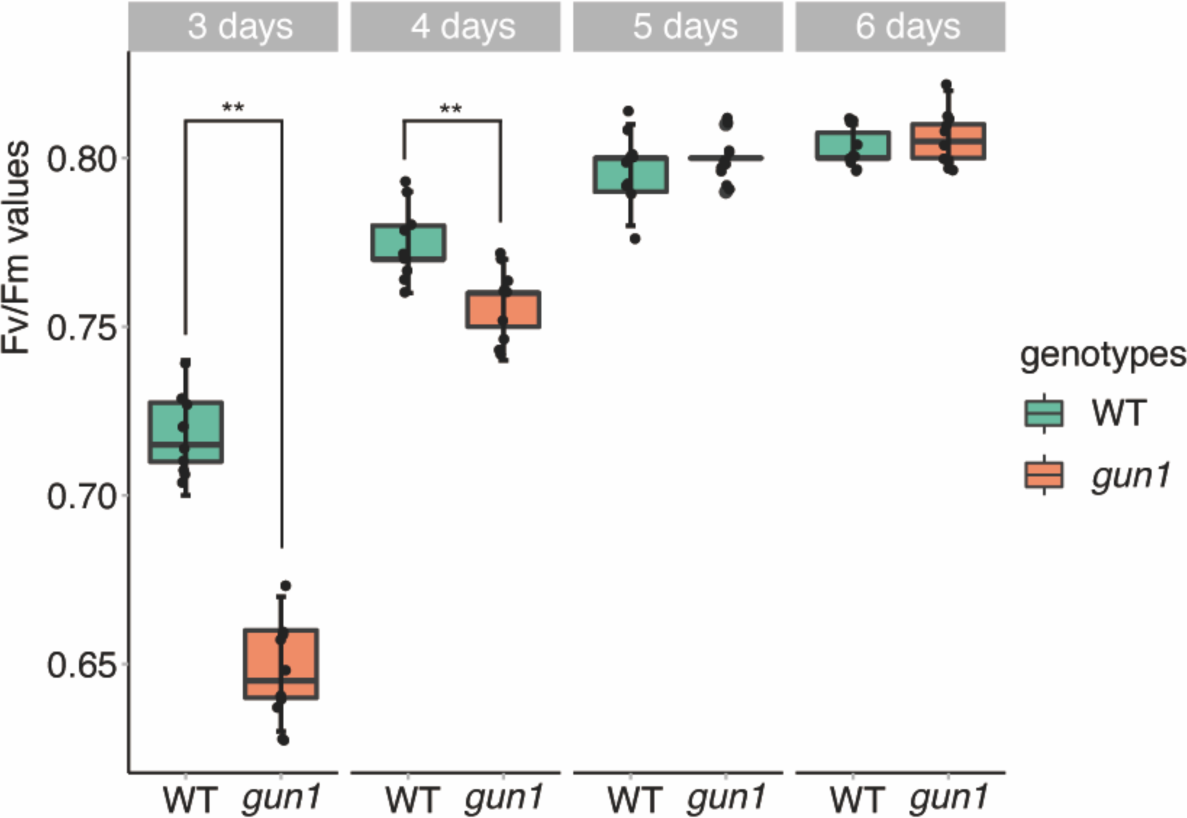
GUN1 promotes PSII biogenesis and photosynthesis during seed germination. F_v_/F_m_ values were determined during seedling development (from 3-to 6-day-old seedlings grown under continuous light conditions). Note that only green seedlings were selected to measure F_v_/F_m_. Values represent the means ± SD (n=10). Stars indicate statistically significant differences between mean values, *P*-values are from two-tailed Student’s *t*-tests. \*\**P* < 0.01.

**Figure S6.**
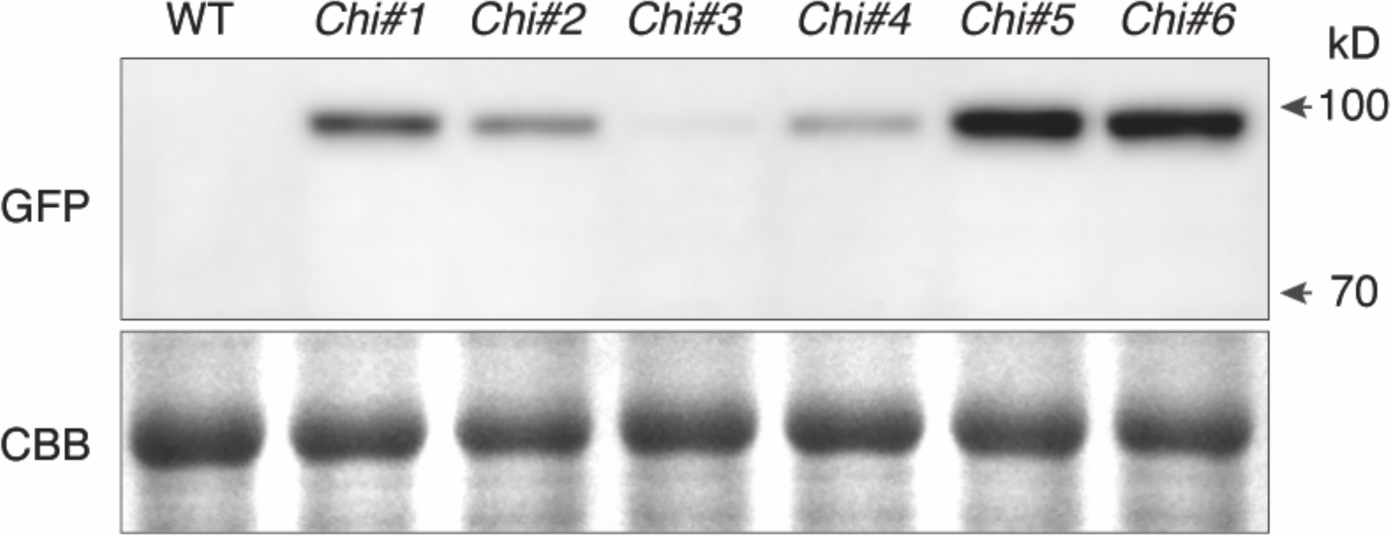
The levels of chimeric CRR4(PPR)-GUN1(SMR) protein fused with GFP in 5-day-old seedlings of individual stable transgenic lines. Immunoblotting assay revealed the steady-state levels of chimeric protein tagged with GFP. Six independent stable homozygous transgenic lines were selected to check the protein expression. *Chi#1* and *Chi#2* were chosen to examine the level of *psbD* BLRP transcripts.

**Table S1.**
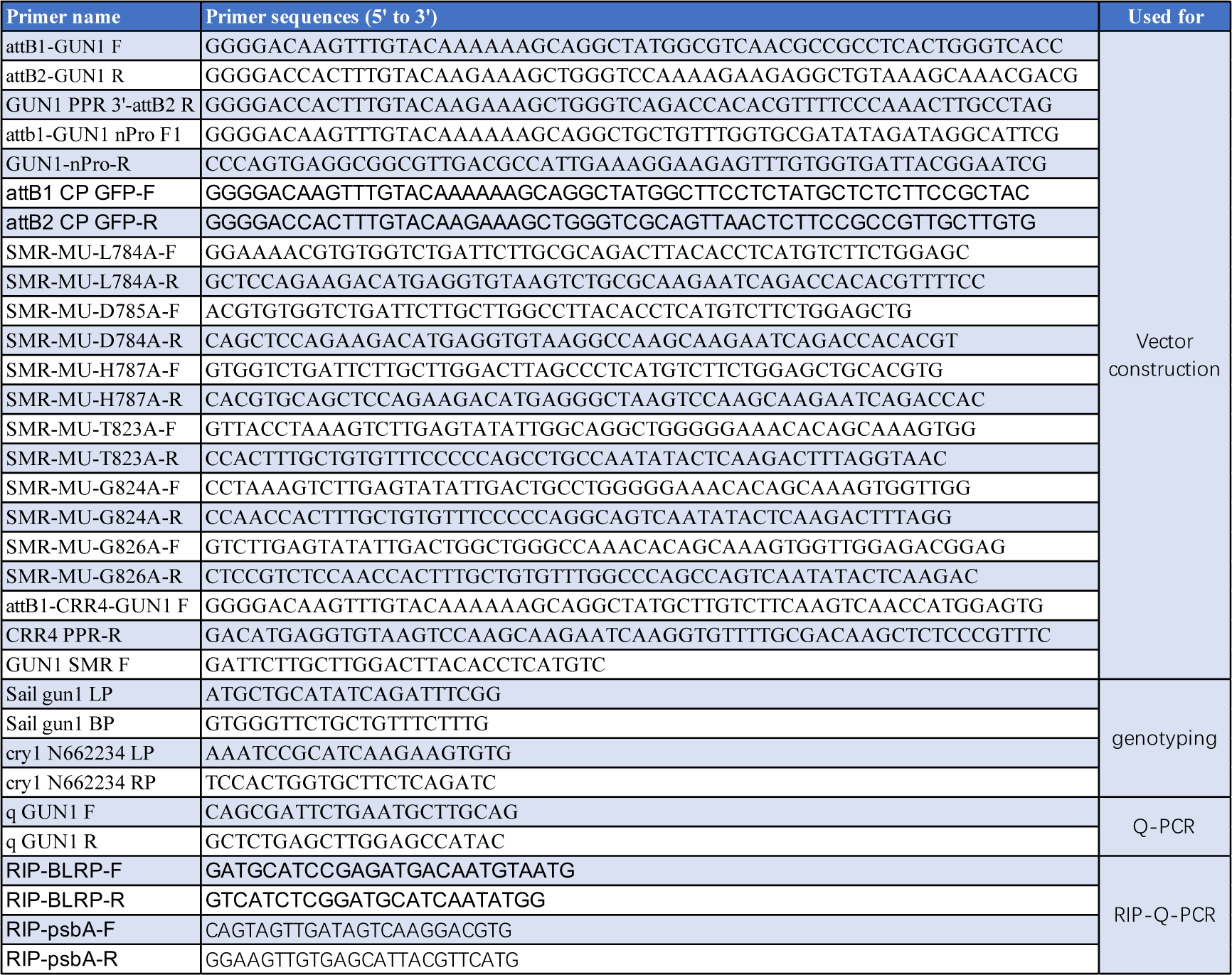
Primers used in this study.

**Table S2.**
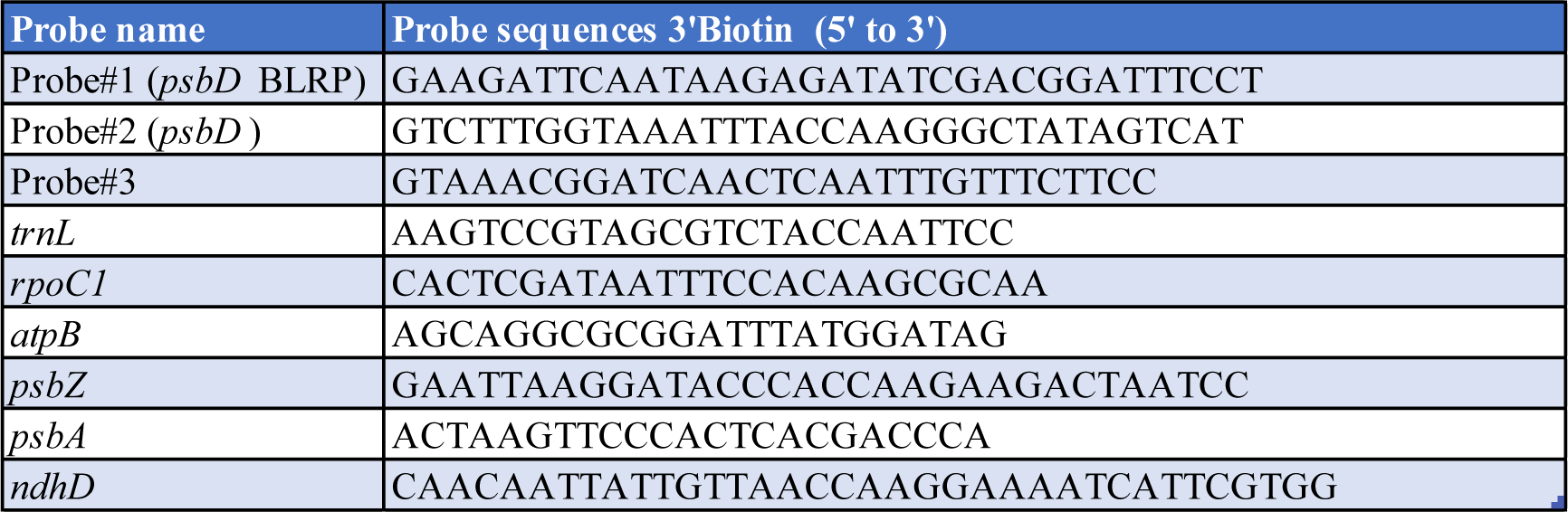
Probes used for Northern Blot assays in this study.

